## Appendix 1 for "Finding structure during incremental speech comprehension"

**Number of figures: 17**

**Number of tables: 3**

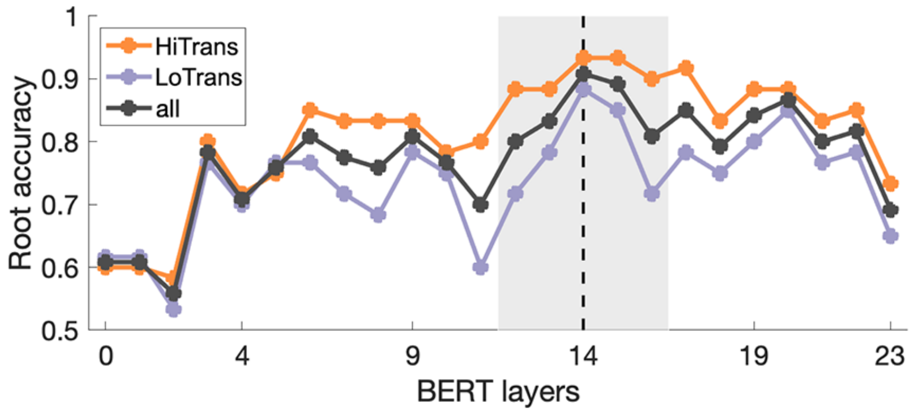


**Appendix 1-figure 1.** Performance of structural probing models trained on different BERT layers. Structural probing models were trained to reconstruct a sentence's structure by estimating each word's dependency parse depth based on the word embeddings from BERT hidden states in this layer. The performance of structural probing models was evaluated by root accuracy (i.e., the percentage of the sentences in which the smallest parse depth is assigned to the main verb, that is, the root of the dependency parse tree which is 0 in the context-free dependency parse tree) when the whole sentence is input to the model. Structural probing models derived from BERT layer 14 showed the best overall performance. However, additional structural probing models derived from its neighboring layers (i.e., layers 12-13, 15-16 in the grey shade) were also included in further analyses to cover potentially useful down-stream and up-stream information. HiTrans: High Verb1 transitivity sentences (n = 60), LoTrans: Low Verb1 transitivity sentences (n = 60), all: HiTrans and LoTrans sentences (n = 120).

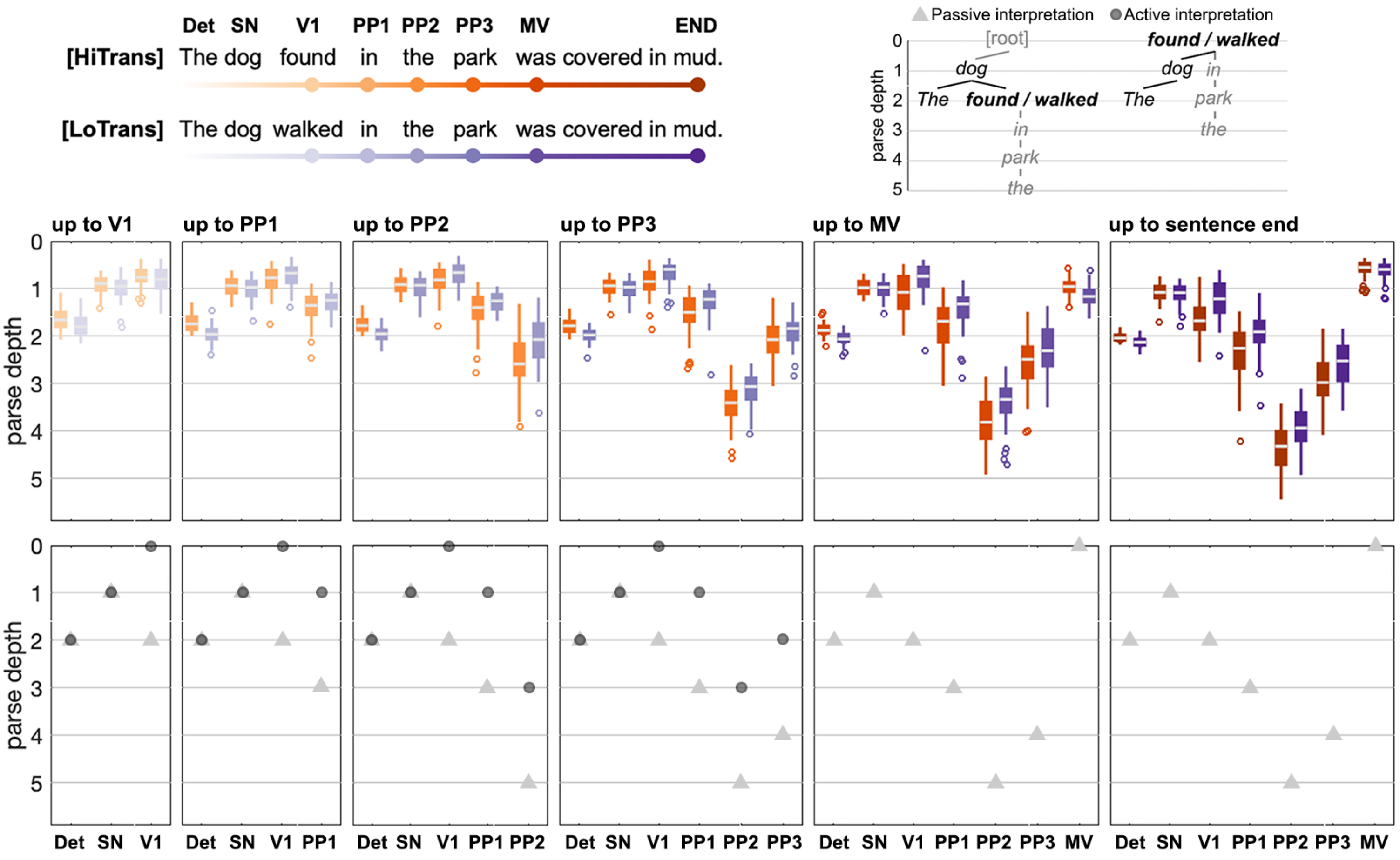


**Appendix 1-figure 2.** BERT structural representations of incremental sentence inputs. Each sentence was input word-by-word to the BERT structural probing model, which returned the estimated parse depth of each word given the specific contents in the incremental sentence input (relevant to **Figures 3D & 3E**). BERT parse depth of words at the same position formed a distribution in both HiTrans and LoTrans sentences (i.e., the boxplots), ranging around the corresponding context-free parse depths in either Passive or Active interpretations denoted by the triangle and circle markers separately, which might reflect probabilistic interpretations given the specific contents in a sentence. Moreover, the BERT parse depth of earlier words in the sentence was updated with each incoming later word, capturing the incrementality of speech comprehension. The results in this figure were obtained from the structural probing model trained on BERT layer 14. Det: determiner, SN: subject noun, V1: Verb1, PP1-PP3: prepositional phrase, MV: main verb, END: end of the sentence.

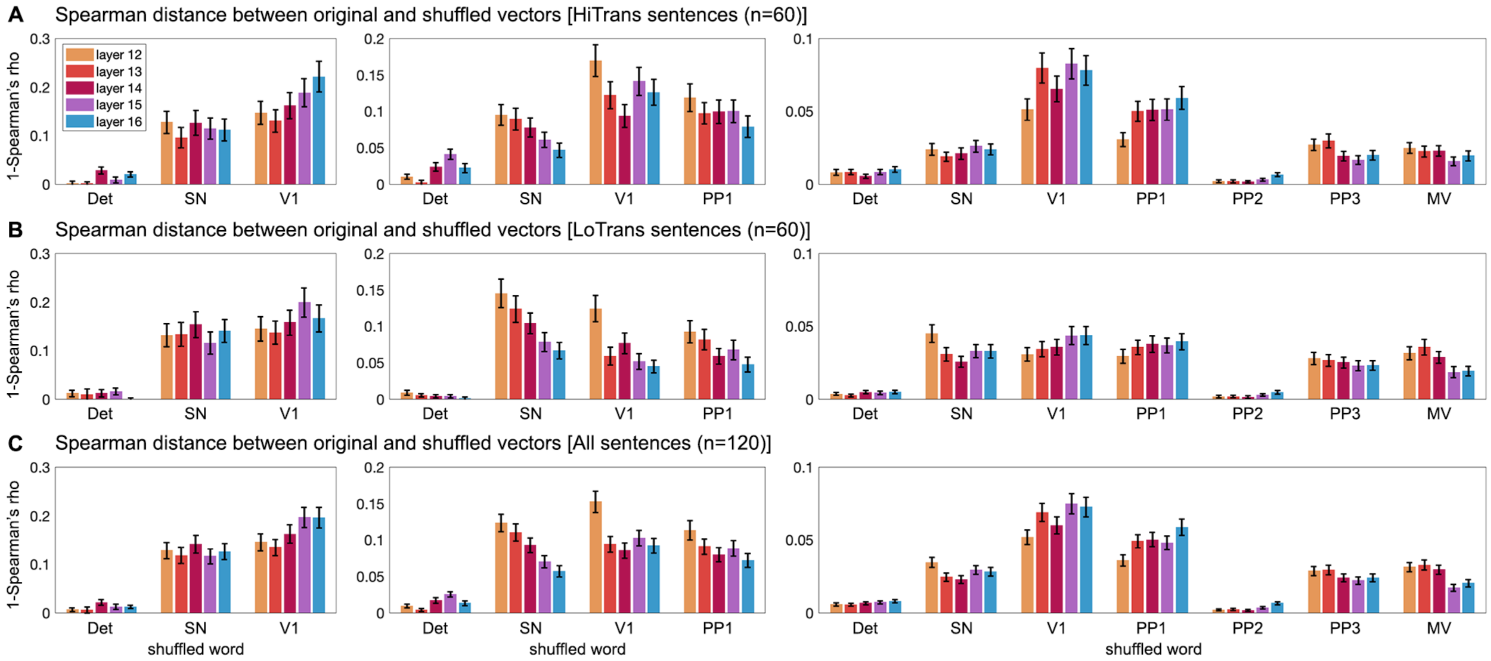


**Appendix 1-figure 3.** Contribution of words at different positions in BERT parse depth vectors (relevant to **Figures 3D & 3E** and **Appendix 1-figure 2**). Given the incremental BERT parse depth vectors up to Verb1 (V1) (left), PP1 (middle) and MV (left), we separately shuffled the parse depths of words at a particular position at a time across **(A)** HiTrans sentences, **(B)** LoTrans sentences or **(C)** both types of sentences, and meanwhile kept the parse depths of the other words unchanged. Then we calculated the Spearman distance (i.e., 1-Spearman's rho) between the original and the shuffled BERT parse depth vectors. The larger this distance, the more the word at this position contributed to the BERT parse depth vector. In general, content words contributed more than function words (i.e., the two determiners, one at the beginning of the sentence (e.g., "***The*** *dog*..."), the other in the prepositional phrase - PP2 (e.g., "*...in* ***the*** *park*..."). In fact, function words contributed the least to the overall variance of BERT structural representations. This analysis was conducted for the BERT parse depth obtained from layers 12-16. BERT parse depth at each position was shuffled 10,000 times, error bars represent one SD. HiTrans: high Verb1 transitivity, LoTrans: low Verb1 transitivity, Det: determiner, SN: subject noun, PP1-PP3: prepositional phrase, MV: main verb.

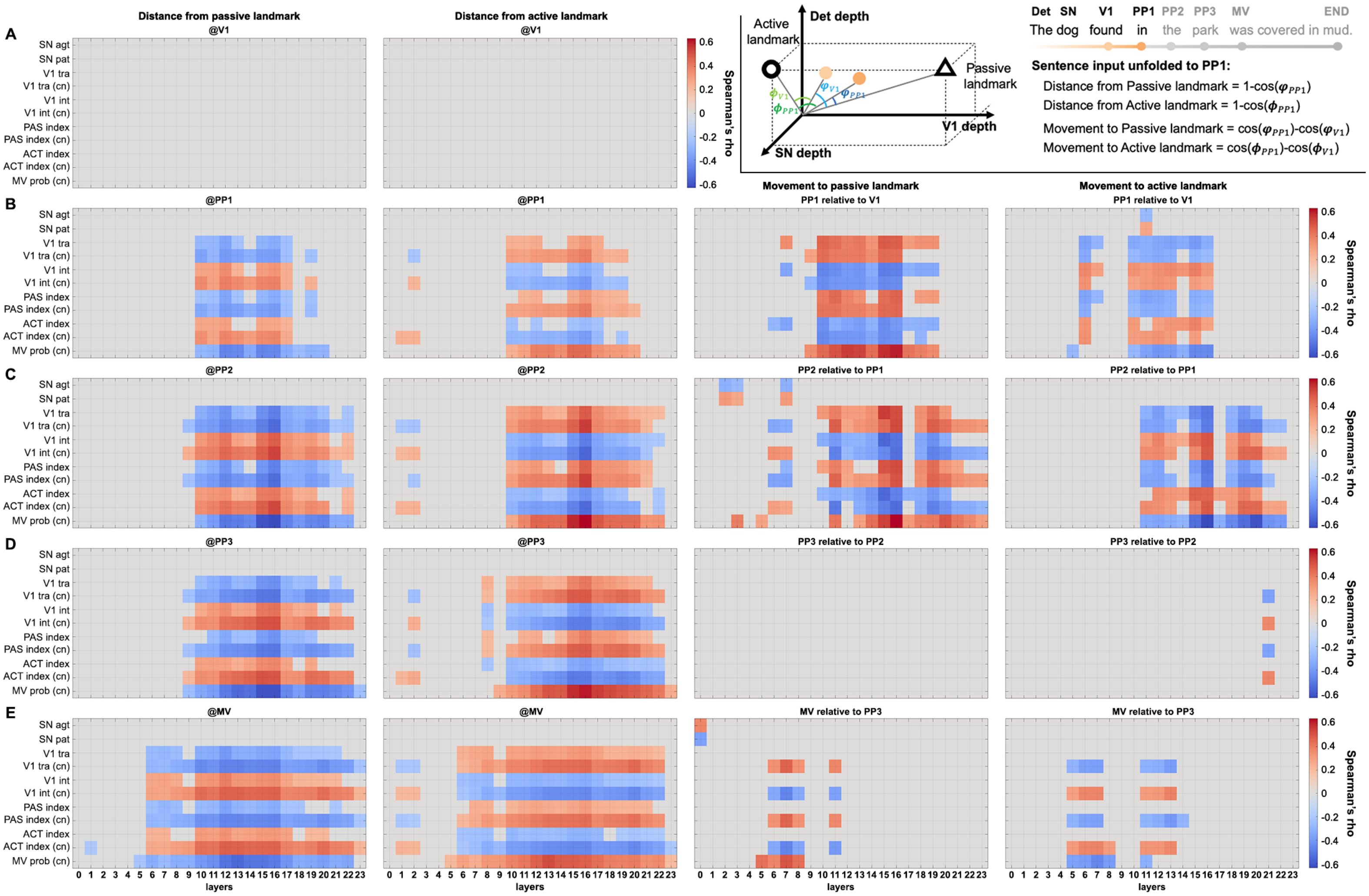


**Appendix 1-figure 4.** Correlation between BERT structural interpretations and explanatory variables (relevant to **Figure 4**). The dynamic interpretation of the structural dependency between the subject noun (SN) and the Verb1 (V1) by BERT can be captured by each sentence's trajectory in BERT model space (see **Figure 3D**), with the Passive/Active landmarks as references. The cosine distance between each sentence and the Passive/Active landmark was calculated as the sentence unfolded word-by-word. The change of the cosine distance from a landmark between two consecutive words (i.e., incremental movement relative to a certain landmark) was also calculated (see an example for a sentence unfolding from V1 to PP1 at the top right corner). The distance and the incremental movement relative to the two landmarks in the model space were correlated with explanatory variables derived from corpus and human continuation pre-tests, correlation results with respect to incremental sentence inputs up to **(A)** V1, **(B)** PP1, **(C)** PP2, **(D)** PP3 and **(E)** MV are separately shown above (Spearman correlation, significance was determined by 10,000 permutations, $P_{FDR} < 0.05$, multiple comparisons corrected for all BERT layers). Note that only significant results are plotted, gray color indicates non-significant results, which also applies to **Appendix 1-figure 5** and **Appendix 1-figure 6**. SN agt/pat: subject noun agenthood/patienthood, V1 tra/int: Verb1 transitivity/intransitivity, PAS/ACT index: Passive/active index, MV prob: probability of a main verb in the continuations after PP derived from the continuation pre-test. Explanatory variables based on human continuations are indicated by "cn" in parentheses. Det: determiner, PP1-PP3: prepositional phrase, MV: main verb.

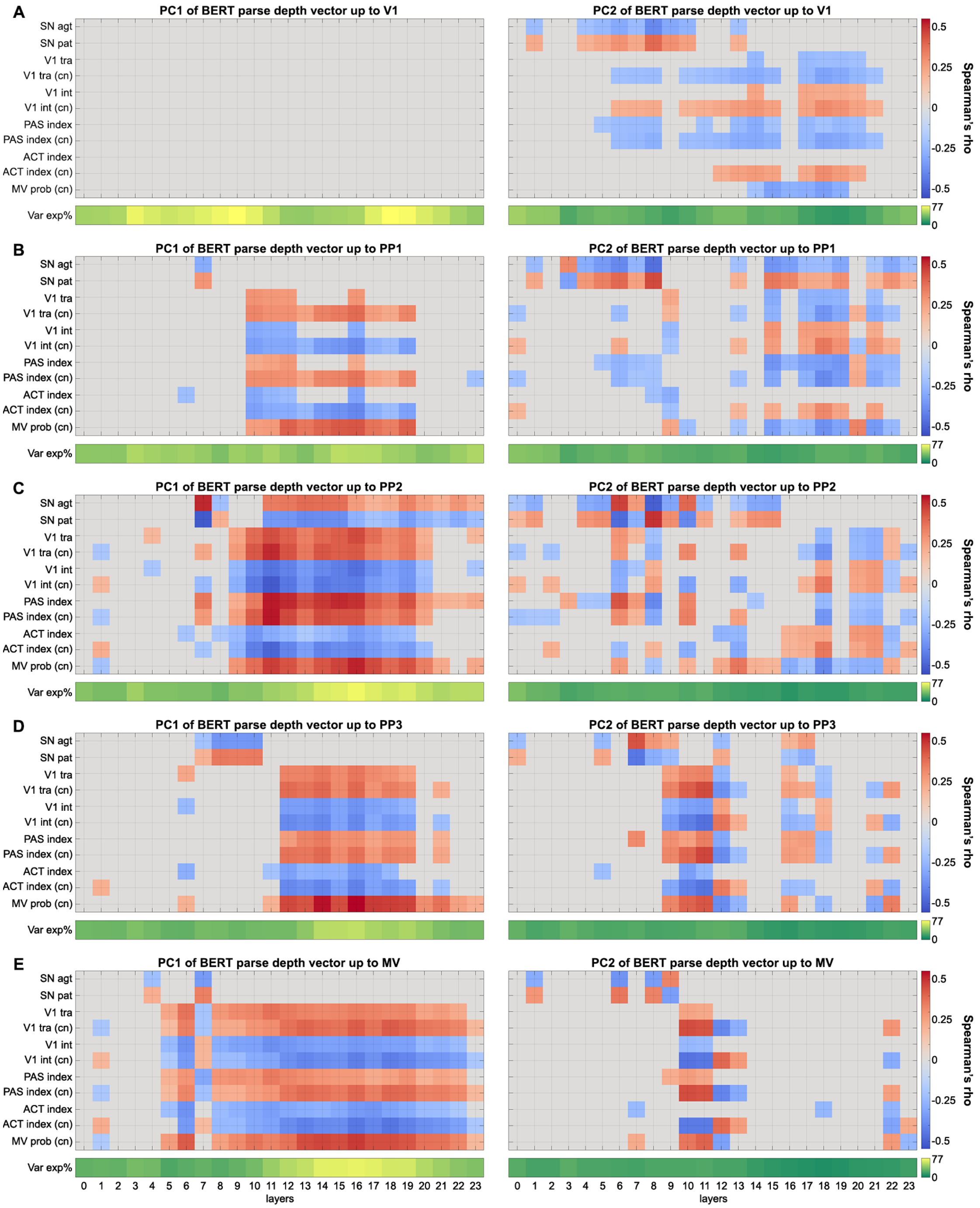


**Appendix 1-figure 5.** Correlation between the principal components (PCs) of BERT parse depth vectors and explanatory variables (relevant to **Figure 4**). PCs of the incremental BERT parse depth vectors were derived to represent the overall information encoded in them. The first two PCs of BERT parse depth vectors for incremental sentence inputs up to **(A)** Verb1 (V1), **(B)** PP1, **(C)** PP2, **(D)** PP3 and **(E)** MV were separately correlated with explanatory variables derived from corpus and human continuation pre-tests (Spearman correlation, significance was determined by 10,000 permutations, $P_{FDR} < 0.05$, multiple comparisons corrected for all BERT layers). The variance explained (Var exp%) by each PC of each BERT layer is shown at the bottom of each panel. Det: determiner, SN: subject noun, PP1-PP3: prepositional phrase, MV: main verb.

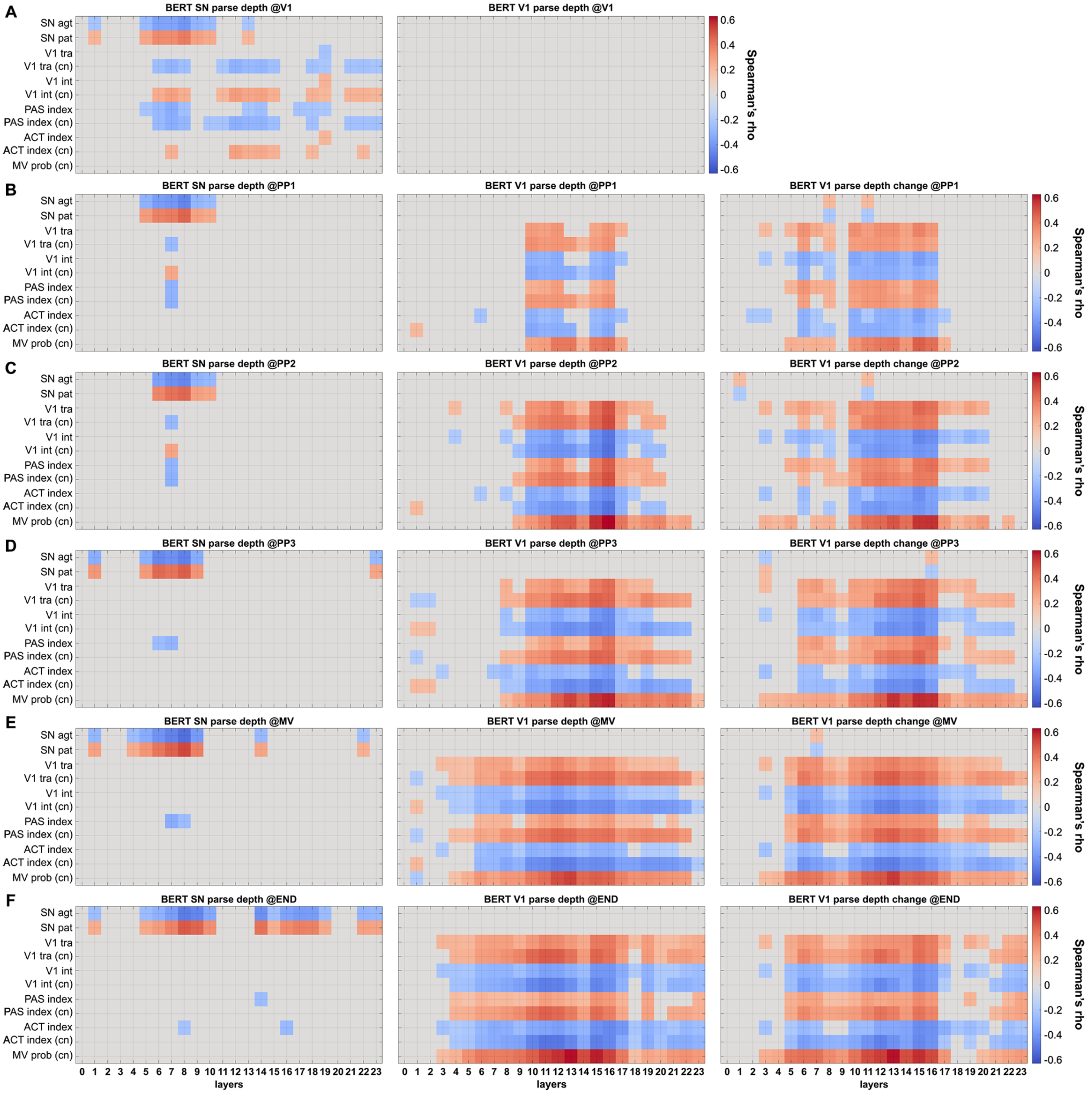


**Appendix 1-figure 6.** Correlation between BERT parse depth of individual words and explanatory variables (relevant to **Figure 4**). BERT parse depth of SN (left), Verb1 (V1) (middle), and BERT Verb1 parse depth change relative to its first appearance (right) obtained from incremental sentence inputs up to **(A)** V1, **(B)** PP1, **(C)** PP2, **(D)** PP3, **(E)** MV and **(F)** the end of the sentence were separately correlated with explanatory variables derived from corpus and human continuation pre-tests (Spearman correlation, significance was determined by 10,000 permutations, $P_{FDR} < 0.05$, multiple comparisons corrected for all BERT layers). Det: determiner, SN: subject noun, PP1-PP3: prepositional phrase, MV: main verb, END: end of the sentence.

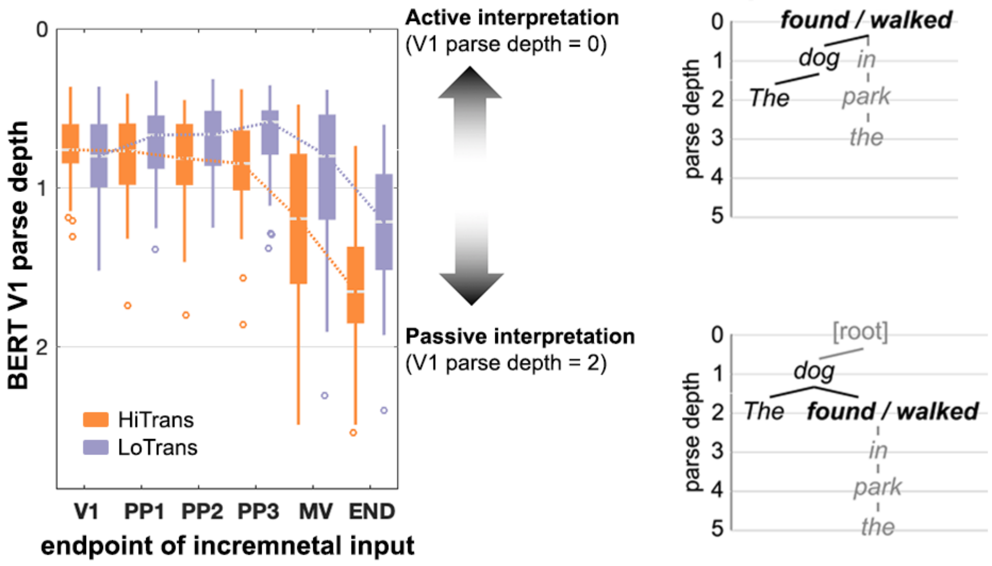


**Appendix 1-figure 7.** The dynamic change of BERT Verb1 (V1) parse depth in unfolding sentences. BERT parse depth of Verb1 was updated every time an incoming later word was added in the incremental sentence input (relevant to **Figure 7**). In the context-free dependency parse tree (right panel), Verb1 parse depth is 2 for a Passive interpretation and is 0 for an Active interpretation. Therefore, the dynamically increasing or decreasing BERT Verb1 parse depth in an unfolding sentence reflected the preference biased towards a Passive or an Active interpretation separately (left panel). In terms of the group-level effect indicated the median value, BERT Verb1 parse depth in HiTrans sentences unidirectionally increased towards the Passive interpretation after Verb1 (i.e., a parse depth of 2), whereas that in LoTrans sentences initially tended to decrease and approached a depth of 0 but increased with the appearance of the actual main verb, suggesting a reorientated preference for the Passive interpretation instead of the initial Active interpretation. The results above were obtained from the structural probing model trained on BERT layer 14. HiTrans: high Verb1 transitivity sentences, LoTrans: low Verb1 transitivity sentences, Det: determiner, SN: subject noun, PP1-PP3: prepositional phrase, MV: main verb, END: end of the sentence.

**V1 epoch** - BERT parse depth vector up to V1 (l13)

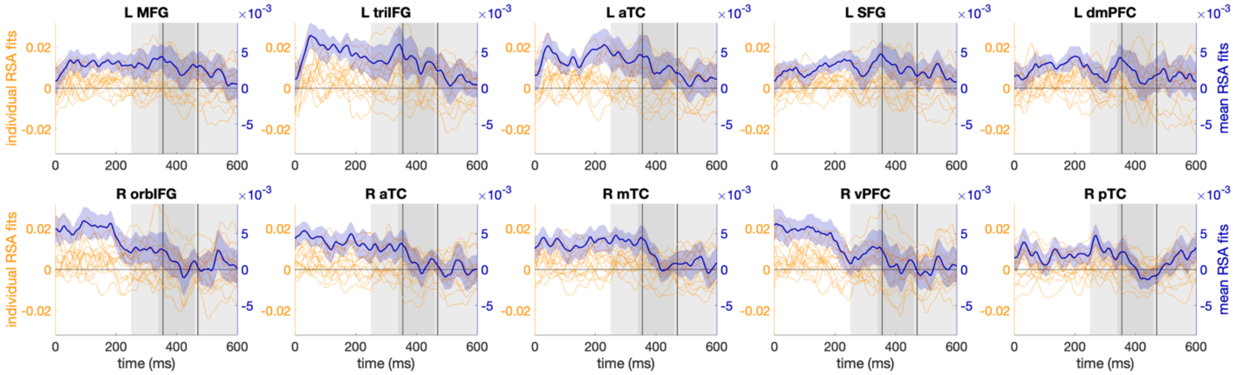


**V1 epoch** - Mismatch for Active interpretation up to V1 (l13)

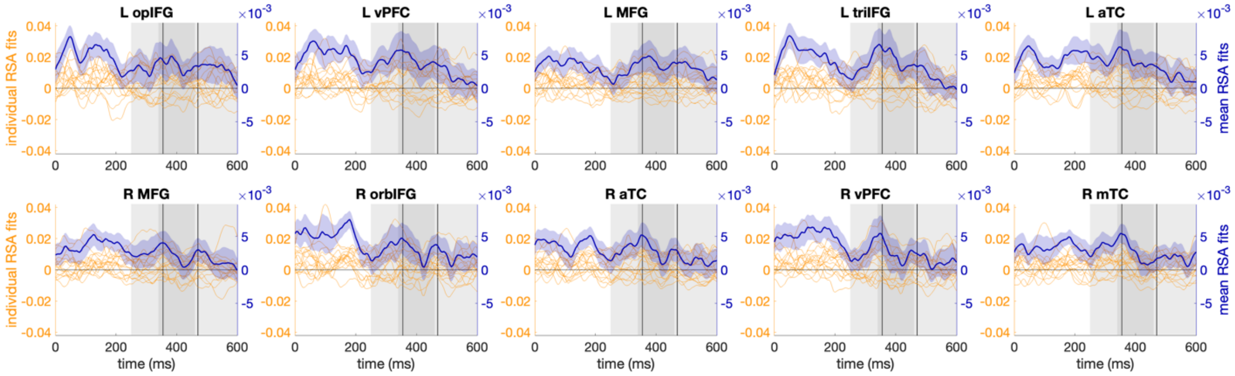


**PP1 epoch** - BERT parse depth vector up to PP1 (l14)

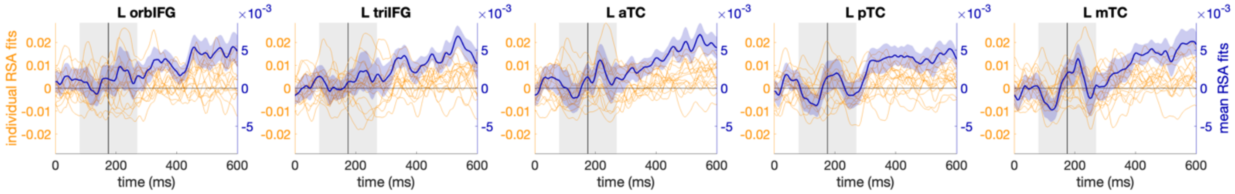


**PP1 epoch** - Mismatch for Passiveinterpretation up to PP1 (l14)

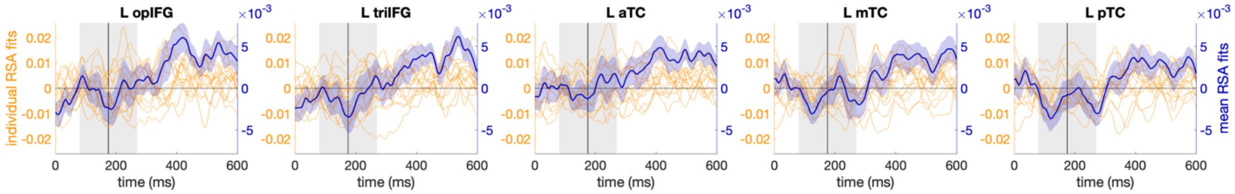


**MV epoch** - BERT parse depth vector up to MV (l14)

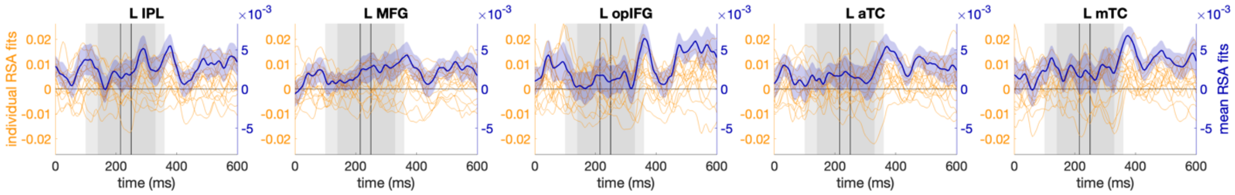


**MV epoch** - Mismatch for Passive interpretation up to MV (l14)

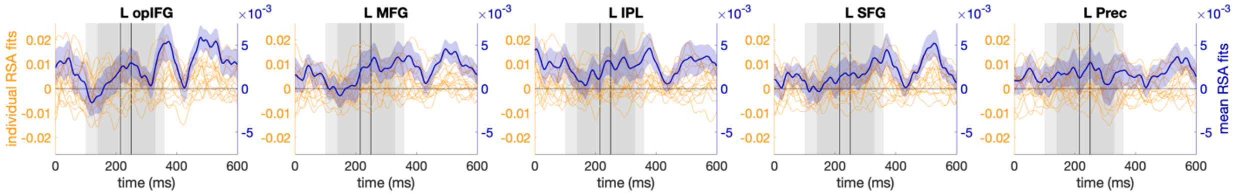

**Appendix 1-figure 8.** Spearman's rho time-series of ROIs across individual participants and their mean (with SEM) for BERT parse depth vector and its mismatch for Active and Passive interpretations in V1, PP1 and MV epochs (relevant to **Figure 6**). V1: Verb1, PP1: preposition, MV: main verb.

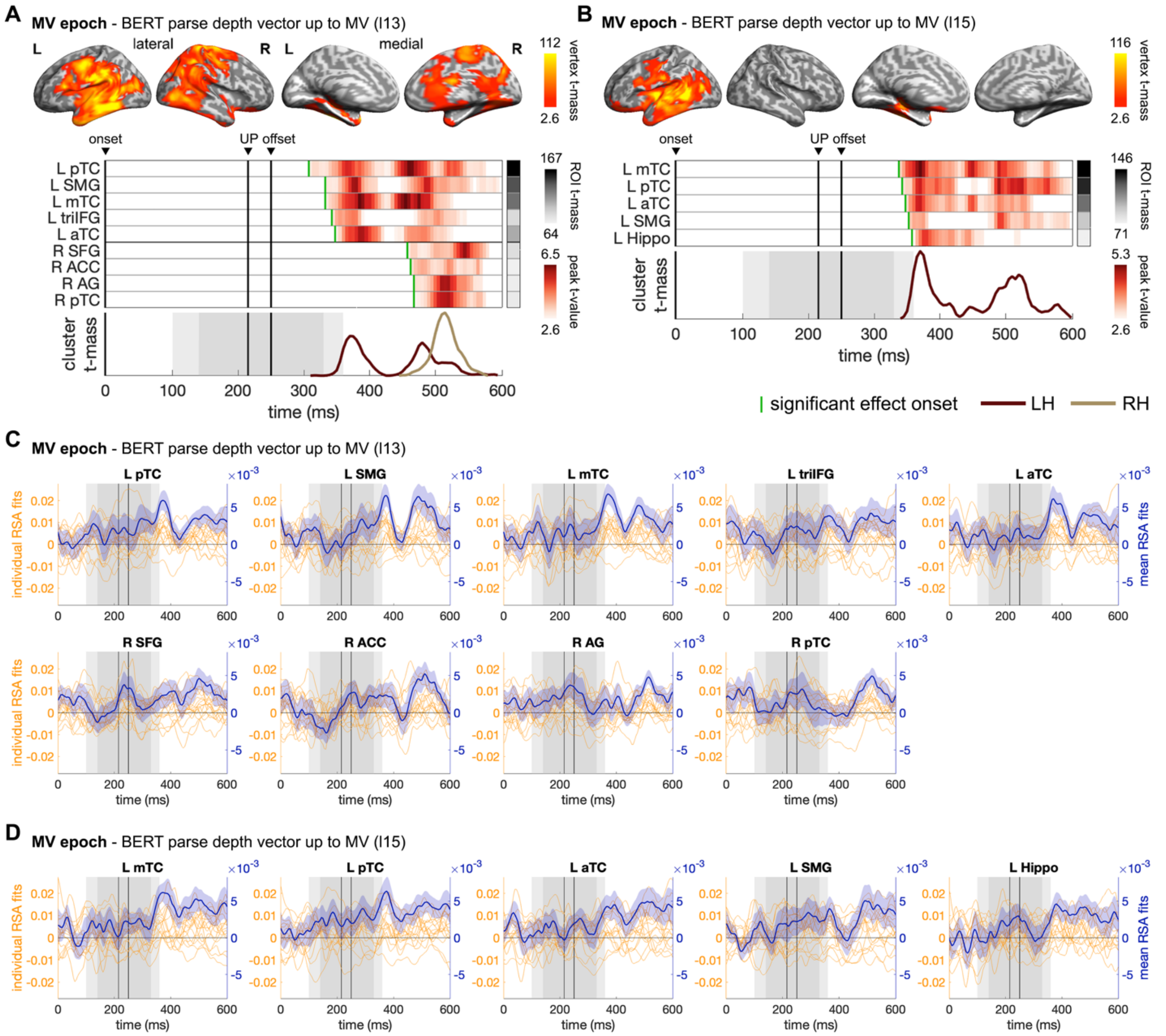


**Appendix 1-figure 9.** ssRSA results of BERT structural measures in the main verb (MV) epoch. ssRSA results of BERT parse depth vector up to MV derived from **(A)** BERT layer 13 and **(B)** BERT layer 15 in MV epoch (cluster-based permutation test, vertex-wise $P < 0.01$, cluster-wise $P < 0.05$). The Spearman's rho time-series of ROIs across individual participants and their mean (with SEM) are shown in **(C)** and **(D)**, respectively.

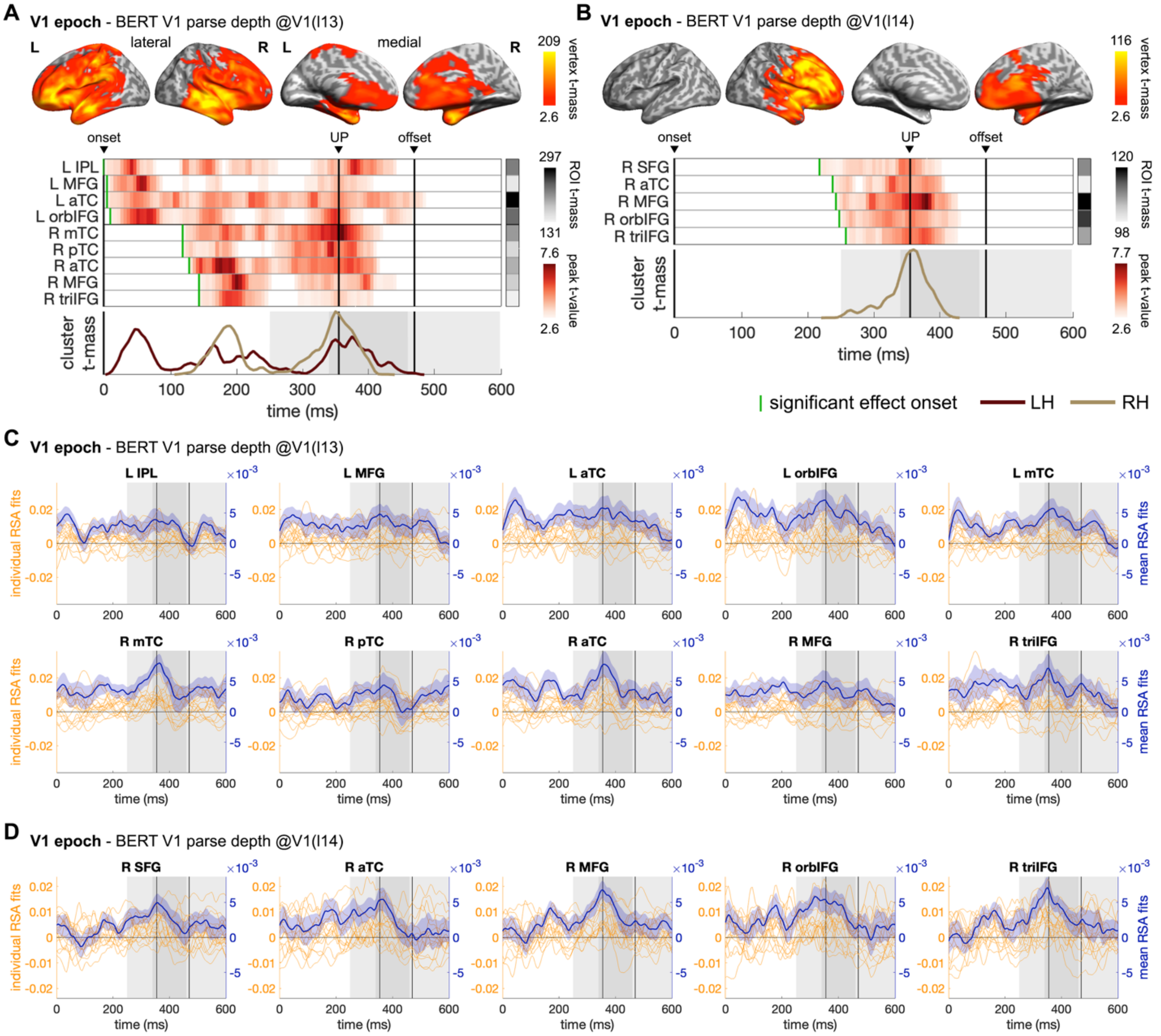


**Appendix 1-figure 10.** ssRSA results of BERT structural measures in the Verb1 (V1) epoch. ssRSA results of BERT Verb1 parse depth derived from **(A)** BERT layer 13 and **(B)** BERT layer 14 in V1 epoch (cluster-based permutation test, vertex-wise $P < 0.01$, cluster-wise $P < 0.05$). From top to bottom in each panel: vertex t-mass (each vertex's summed t-value during its significant period); time-series of ROI peak t-value (the highest t-value in an ROI at each time-point with a green bar indicating effect onset) and ROI t-mass (each ROI's summed mean t-value during its significant period); cluster t-mass time-series (summed t-value of all the significant vertices of a cluster at each time-point). Solid vertical lines indicate the timings of onset, average uniqueness point (UP) and average offset of the word time-locked in the epoch with grey shades indicating the range of one SD. LH/RH: left/right hemisphere. The Spearman's rho time-series of ROIs across individual participants and their mean (with SEM) are shown in **(C)** and **(D)**, respectively.

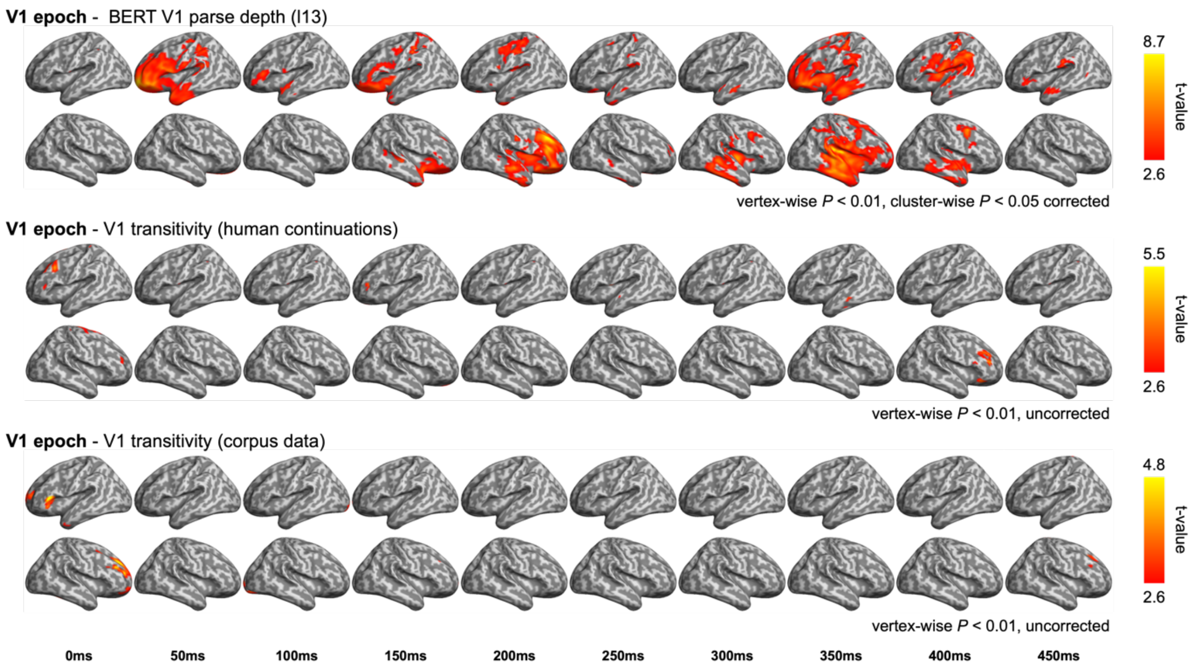


**Appendix 1-figure 11.** Comparison between the RSA model fits of BERT structural metrics and behaviour- / corpus-based metrics in the Verb1 (V1) epoch. (upper) Model fits of BERT Verb1 parse depth (relevant to **Appendix 1-figure 10A**); (middle) Model fits of the first verb transitivity based on the continuation pre-rest conducted by the end of V1 (e.g., complete "*The dog found* …"); (bottom) Model fits of the Verb1 transitivity based on the same corpus data described in Methods.

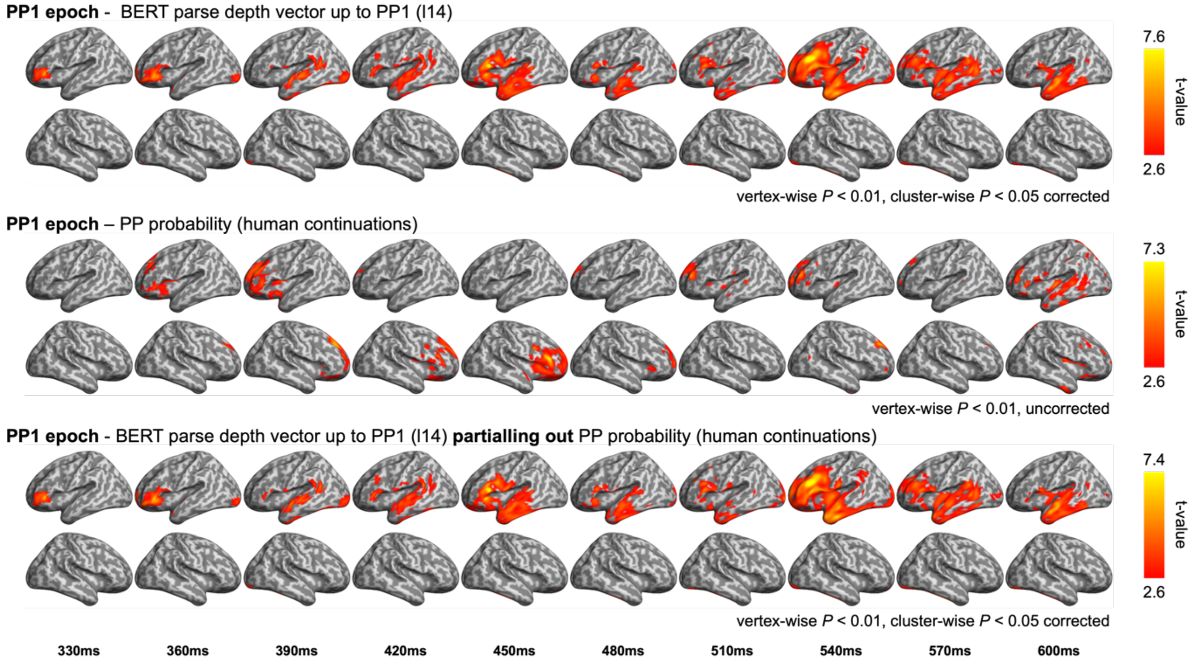


**Appendix 1-figure 12.** Comparison between the RSA model fits of BERT structural metrics and behaviour- / corpus-based metrics in the preposition (PP1) epoch. (upper) Model fits of BERT parse depth vector up to PP1 (relevant to **Figure 6B**); (middle) Model fits of the probability of a prepositional phrase (PP) continuation in the pre-rest conducted by the end of the first verb (e.g., complete "*The dog found* …"); (bottom) Model fits of BERT parse depth vector up to PP1 while partialling out the variance explained by PP probability.

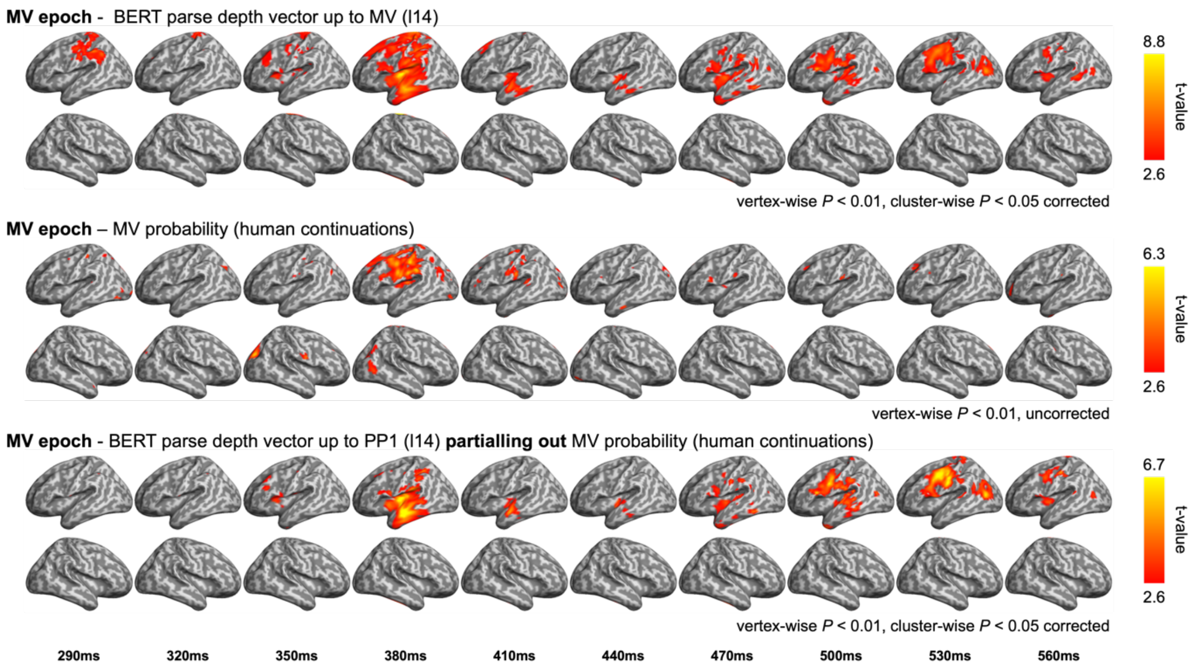


**Appendix 1-figure 13.** Comparison between the RSA model fits of BERT structural metrics and behaviour- / corpus-based metrics in the main verb (MV) epoch. (upper) Model fits of BERT parse depth vector up to MV (relevant to **Figure 6C**); (middle) Model fits of the probability of a main verb continuation in the pre-rest conducted by the end of the prepositional phrase (e.g., "*The dog found in the park* …"); (bottom) Model fits of BERT parse depth vector up to MV while partialling out the variance explained by MV probability.

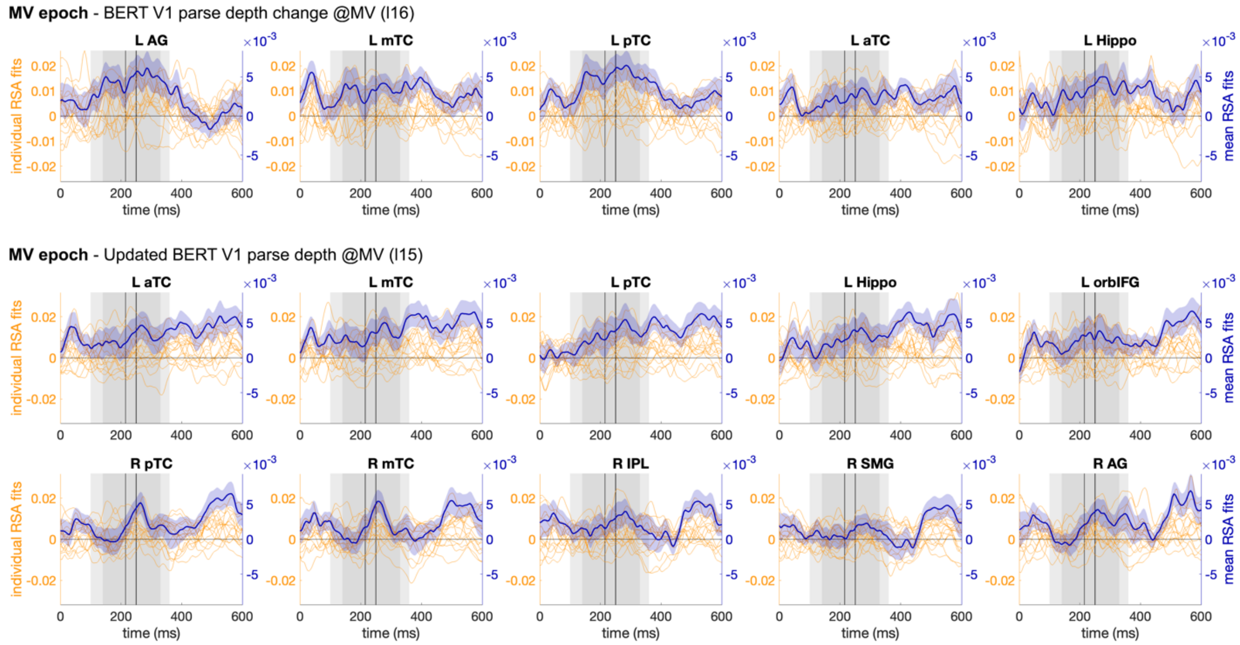


**Appendix 1-figure 14.** Spearman's rho time-series of ROIs across individual participants and their mean (with SEM) for BERT V1 parse depth change and the updated BERT V1 parse depth in the MV epoch (relevant to **Figure 7**). V1: Verb1, MV: main verb.

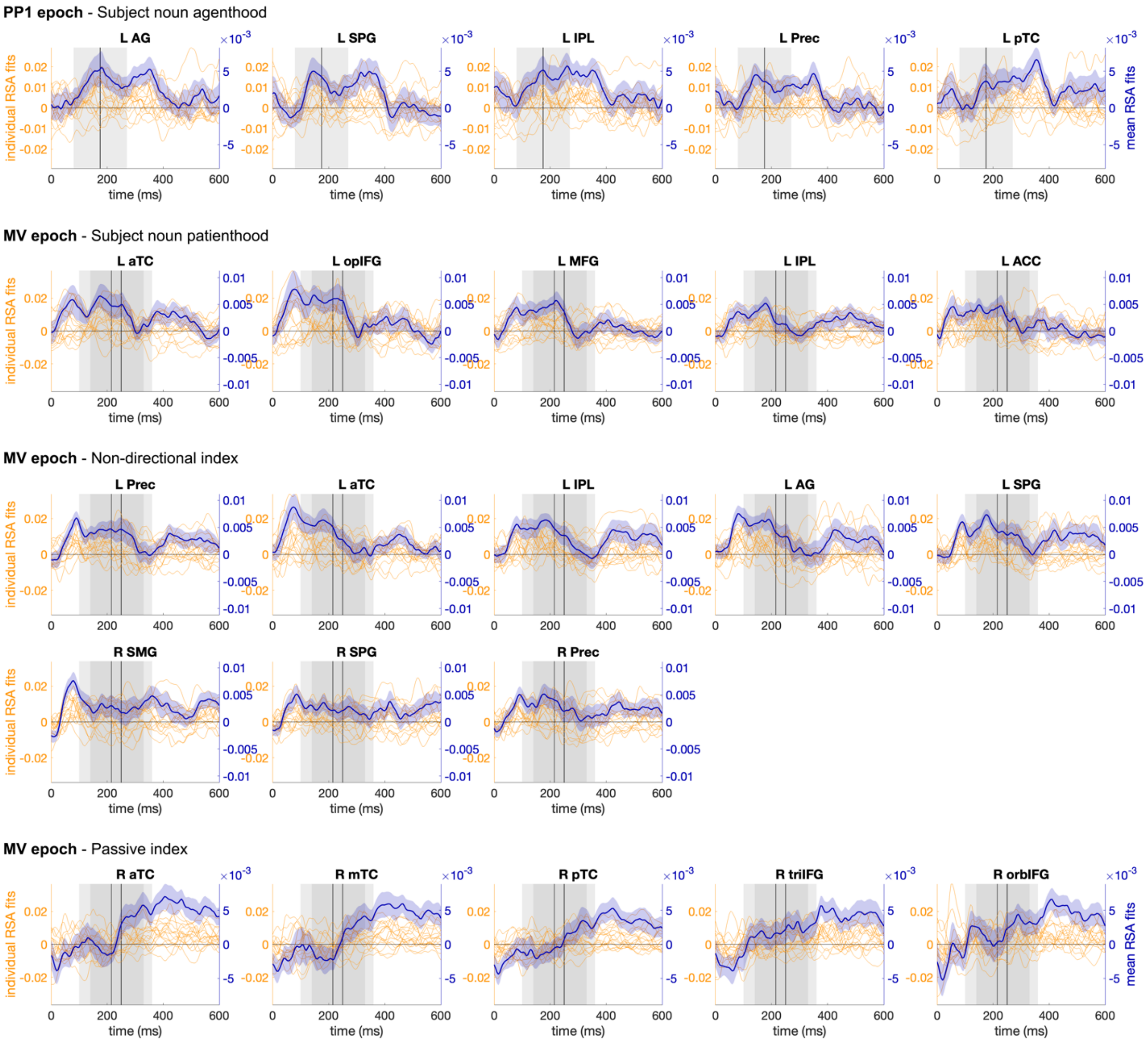


**Appendix 1-figure 15.** Spearman's rho time-series of ROIs across individual participants and their mean (with SEM) for corpus-based measures in PP1 and MV epochs (relevant to **Figure 8**). PP1: preposition, MV: main verb.

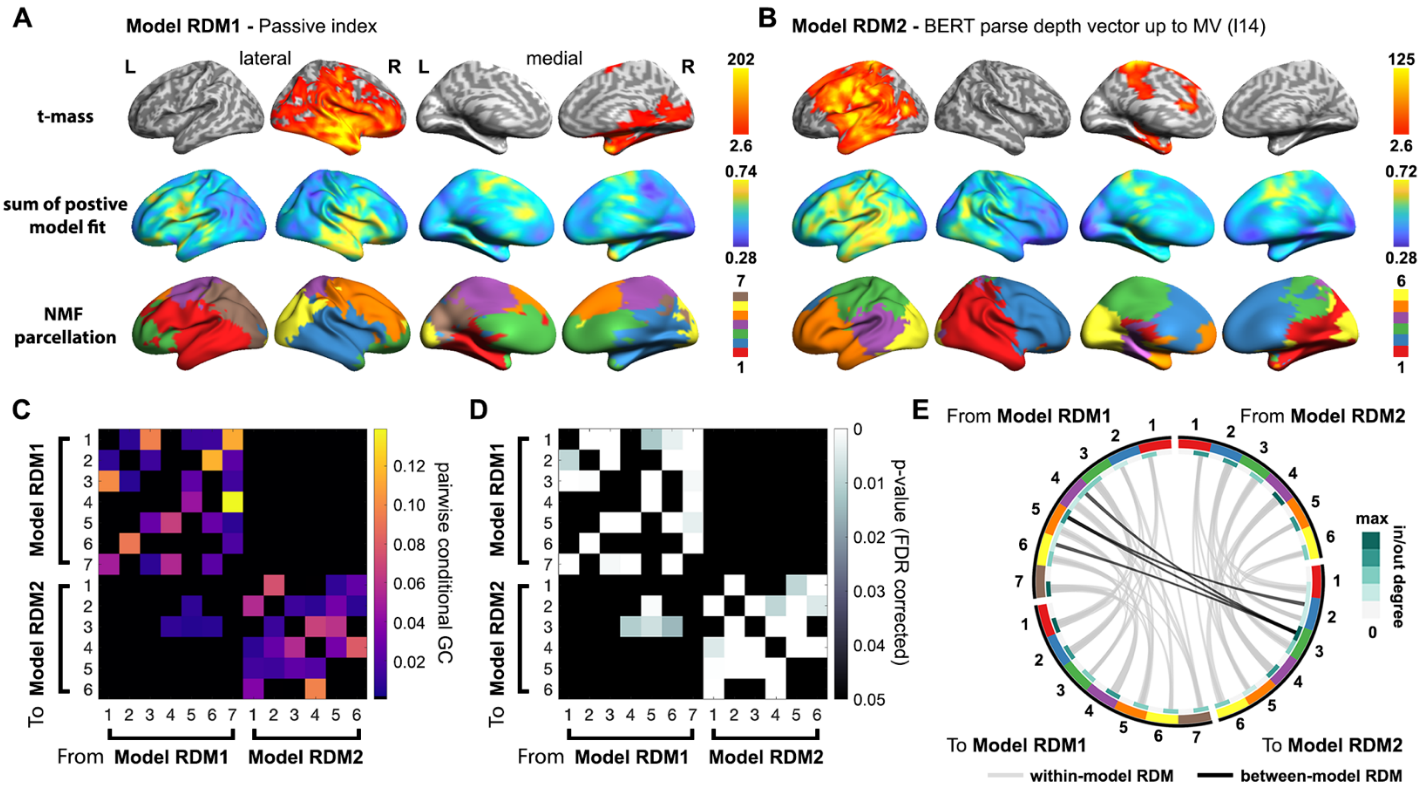


**Appendix 1-figure 16.** Directional relationship between multifaceted constraints and structured interpretation in the brain (relevant to **Figure 8E** in the main text). Non-negative matrix factorization (NMF) was applied to the whole-brain ssRSA models fits of Passive index and BERT parse depth vector up to the main verb (MV) from layer 14 in the MV epoch separately. Multivariate Granger causality analysis (GCA) was conducted based on time-series of the NMF factors of the two model RDMs. Vertex-wise results of ssRSA and NMF parcellation of **(A)** Passive index and **(B)** BERT parse depth vector up to MV: (from top to bottom) each vertex's summed t-value during its significant period, sum of uncorrected positive model fit averaged across participants, NMF parcellation results illustrated by assigning each vertex to the NMF factor with the highest loading value. **(C)** Pairwise conditional GC and **(D)** corresponding p-values obtained from multivariate GCA based on the two sets of NMF factors (FDR corrected $P < 0.05$). **(E)** Circos plot of significant GC connections between model RDMs in (C) and (D).

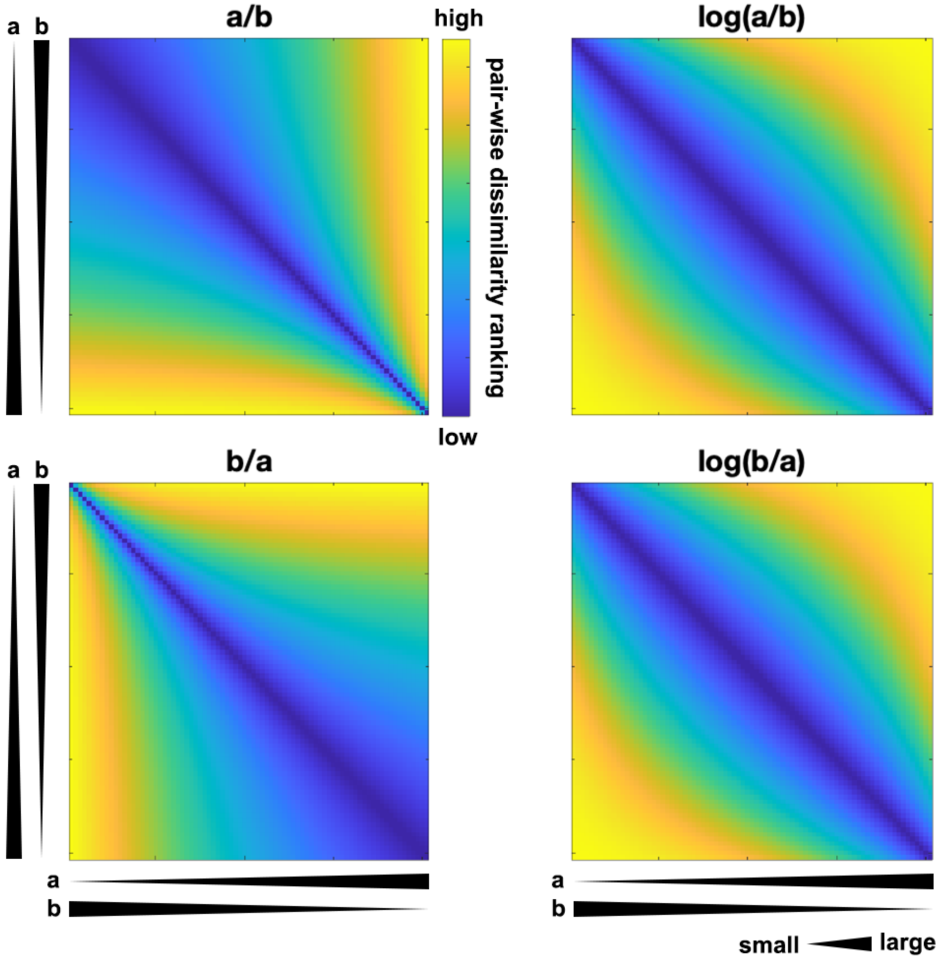


**Appendix 1-figure 17.** Illustration of directionality of dissimilarity geometry in the representational dissimilarity matrix (RDM) based on a ratio measure. Suppose that variable ***a*** ranges from 0.1 to 0.9, and there are 81 items evenly sampled with an increment of 0.01 from ***a***, while variable ***b*** equals 1-***a***. Items are sorted by their values as indicated by the black wedges corresponding to ***a*** and ***b*** (ascending order for ***a***, descending for ***b*** from top to bottom, from left to right). RDMs are built by calculating the absolute pairwise difference of ***a/b***, ***b/a*** (left) and their logarithmic transformation counterparts log(***a/b***) and log(***b/a***) (right). RDMs of ***a***/***b*** or ***b***/***a*** are directional in the sense that they distinguish items with large values for the numerator from the other items in the dissimilarity geometry [i.e., the contrast between the bright yellow (larger dissimilarity) and dark blue (smaller dissimilarity) areas in the RDM]. However, such directionality is removed once the logarithmic transformation is applied, given that |log(***a***)-log(***b***)| is the same as |log(***b***)-log(***a***)|.

**Appendix 1-table 1.** Summary of all quantitative measures used to create model RDMs

| Name | Type | Description | Results |
|---|---|---|---|
| BERT parse depth vector | vector | a N-dimensional vector consisting of the BERT parse depths of all the N words in an incremental input. i.e., BERT's structural representation of this incremental input | Figs. 6A-6C<br><br>Appx1. Figs. 8, 9, 12, 13, 16 |
| BERT interpretative mismatch | scalar | cosine distance between a BERT parse depth vector and the corresponding context-free parse depth vector for the Passive or the Active interpretation given the same incremental sentence input (the smaller the mismatch with one interpretation, the higher the preference for this interpretation) | Figs. 6D-6F<br><br>Appx1. Fig. 8 |
| BERT Verb1 parse depth | scalar | BERT parse depth of Verb1 which is updated with each incoming later word, with increased or decreased depth reflecting the preference biased to a passive verb (i.e., Passive interpretation) or the main verb (i.e., Active interpretation) separately | Fig. 7B<br><br>Appx1. Figs. 10, 11, 14 |
| BERT Verb1 parse depth change at MV | scalar | the difference between the updated BERT parse depth of Verb1 when the actual main verb (MV) is heard and its initial value when Verb1 is first encountered, which reflects the efforts required to resolve the potential structural ambiguity | Fig. 7A<br><br>Appx1. Fig. 14 |
| Subject noun agenthood | scalar | the ratio of the number of the subject noun's appearances as an agent to that of its appearances as a patient (estimated from large corpora) | Fig. 8A,<br><br>Appx1. Fig. 15 |
| Subject noun patienthood | scalar | the ratio of the number of the subject noun's appearances as a patient to that of its appearances as an agent (estimated from large corpora) | Fig. 8B,<br><br>Appx1. Fig. 15 |
| Verb1 transitivity | scalar | the ratio of the frequency Verb1 takes a direct object to that it takes | Appx1. Fig. 11(uncorrected) |

| | | | |
|---|---|---|---|
| | | alternative subcategorization frames (SCFs) (estimated from large corpora or human continuation data) | |
| Verb1 intransitivity | scalar | the ratio of the frequency Verb1 takes intransitive SCFs to that it takes a direct object (estimated from large corpora or human continuation data) | No significant results |
| PP probability | scalar | probability of a prepositional phrase continuation from the results of pre-test conducted by the end of the first verb (e.g., complete "*The dog found …*") | Appx1. Fig. 12 |
| MV probability | scalar | probability of a main verb continuation from the results of pre-test conducted by the end of the prepositional phrase (e.g., complete "*The dog found in the park …*") | Appx1. Fig. 13 |
| Active index | scalar | the product between subject noun agenthood and Verb1 intransitivity, capturing the interpretative coherence between each pair of subject noun and Verb1 for an Active interpretation | No significant results |
| Passive index | scalar | the product between subject noun patienthood and Verb1 transitivity, capturing the interpretative coherence between each pair of subject noun and Verb1 for a Passive interpretation | Fig. 8D<br><br>Appx1. Fig. 16 |
| Non-directional index | scalar | obtained by applying logarithmic transformation to the ratio measures of lexical constraints (i.e., subject noun agenthood or patienthood, Verb1 transitivity or intransitivity) before multiplying them, capturing the interpretative coherence between each pair of subject noun and Verb1 regardless of which interpretation is considered | Fig. 8C |

**Appendix 1-table 2.** Full anatomical labels of abbreviations in Figs. 6, 7, and 8, Appx1. Figs. 9 and 10.

| **Abbreviation** | **Full anatomical label** |
|---|---|
| opIFG | Inferior frontal gyrus, opercular part |
| triIFG | Inferior frontal gyrus, triangular part |
| orbIFG | Inferior frontal gyrus, orbital part |
| SFG | Superior frontal gyrus |
| MFG | Middle frontal gyrus |
| dmPFC | Dorsal medial prefrontal cortex |
| vPFC | Ventromedial prefrontal cortex |
| ACC | Anterior cingulate cortex |
| aTC | Anterior temporal cortex |
| mTC | Middle temporal cortex |
| pTC | Posterior temporal cortex |
| AG | Angular gyrus |
| SMG | Supramarginal gyrus |
| SPG | Superior parietal gyrus |
| IPL | Inferior parietal lobe |
| Prec | Precuneus |
| Hippo | Hippocampus |

**Appendix 1-table 3.** Example stimuli sentence set presented to both BERT and human listeners.

| **Conditions** | | | | | | | | | | |
|---|---|---|---|---|---|---|---|---|---|---|
| **UNA** | The | dog | that was | found | in | the | park | | was | covered in mud. |
| **HiTrans** | The | dog | | found | in | the | park | | was | covered in mud. |
| **LoTrans** | The | dog | | walked | in | the | park | | was | covered in mud. |
| **PAS** | The | dog | was | found | in | the | park | and | was | covered in mud. |
| **DO1** | The | king | | found | | the | coat | | behind | the golden throne. |
| **DO2** | The | king | | walked | | the | queen | | around | the palace garden. |
